# Neutrophil Extracellular Traps Drive a skin–kidney inflammatory axis in UVB-Irradiated Lupus-Prone Mice

**DOI:** 10.1101/2023.12.23.572573

**Authors:** Xing Lyu, Minghui Yi, Minghui Li, Ping L. Zhang, Wei Wei, Victoria P. Werth, Ming-Lin Liu

**Affiliations:** Corporal Michael J. Crescenz VAMC, Philadelphia, PA, USA; Department of Dermatology, Perelman School of Medicine, University of Pennsylvania, Philadelphia, PA, USA; Division of Anatomic Pathology, Beaumont Laboratories, Beaumont Health, Royal Oak, Michigan 48073, USA; Department of Rheumatology and Immunology, Tianjin Medical University General Hospital, Tianjin, China

**Author notes:** **Correspondence to:** Ming-Lin Liu, MBBS, PhD, and Victoria P. Werth, MD, Department of Dermatology, Perelman School of Medicine, University of Pennsylvania, and Philadelphia Veterans Administration Medical center. Room B414, 3900 Woodland Avenue, Philadelphia, PA 19104.

**Keywords:** SLE, NETosis, PKCα, CXCR4, Skin-kidney communication

## Abstract

Neutrophil extracellular traps (NETs) are major pathogenic effectors in chronic systemic lupus erythematosus (SLE) pathogenesis, but how they mediate acute, UVB-triggered skin inflammation and subsequent systemic injury remains poorly defined. Using UVB-irradiated lupus-prone mice, we identified an acute NETosis-driven skin-kidney inflammatory axis linking photosensitivity to renal inflammation. UVB induced concurrent skin and kidney inflammation with neutrophil infiltration, NETosis, and release of NET-associated effector cytokines/complement C3 and proteinuria. Mechanistically, PKCα-dependent nuclear envelope rupture licensed NETosis, whereas *Pkcα* deletion markedly suppressed NET formation and inflammation in both organs. Interestingly, only a subset of skin-infiltrating neutrophils underwent local NETosis. Spatiotemporal photoconversion tracking revealed that surviving, CXCR4-expressing neutrophils disseminated from UVB-irradiated skin to kidneys, where they underwent secondary NETosis, delivering cytokines/C3 to drive renal inflammation. Genetic or pharmacological CXCR4 inhibition blocked this dissemination, attenuating renal inflammation and proteinuria. These findings identify an acute NETosis-driven skin-kidney axis, highlighting PKCα-dependent NETosis and CXCR4-mediated neutrophil dissemination as therapeutic targets in photosensitive lupus.

## INTRODUCTION

Systemic lupus erythematosus (SLE) is a systemic autoimmune disease characterized by inflammatory damage to multiple organs, such as the skin and kidneys. The pathogenesis of SLE involves immune cells, cytokines, immune complexes (ICs), and complement, which drives inflammation and tissue damage (*1*). While the etiology remains a complex interplay of genetic, epigenetic, and environmental factors (*1, 2*), the discordance of SLE in monozygotic twins highlights the crucial role of environmental triggers (*1*).

Sunlight has been recognized as a primary trigger for lupus flares since the 1960s (*3*). Over 70% of cutaneous lupus patients are photosensitive (*4*), and ultraviolet B (UVB) overexposure exacerbates both local skin inflammation and systemic symptoms, including lupus nephritis, in SLE patients and autoimmune mice (*5–8*). This link is supported by epidemiological data showing a seasonal pattern for severe manifestations, like lupus nephritis, in high-sunlight regions (*9, 10*). However, the mechanisms linking cutaneous UVB irradiation to distant organ injury remain a critical knowledge gap.

Neutrophils are the first circulating leukocytes to infiltrate the skin following UVB overexposure (*11*) and act as initiator immune cells driving early spontaneous CLE lesions in a lupus mouse model (*12*). We previously demonstrated that UVB irradiation triggers skin inflammation and neutrophil infiltration, with a fraction of these cells undergoing NETosis―the release of neutrophil extracellular traps (NETs) (*13, 14*). While NETosis has been implicated in the chronic pathogenesis of cutaneous lupus (*15, 16*) and lupus nephritis (*17, 18*), whether NETosis mechanistically initiates acute systemic disease exacerbation following UVB-induced photosensitivity remains unknown.

NETs are potent inducers of type I interferon (IFN-I) (*14, 19*), a primary driver of lupus pathogenesis (*20*). Together, NETs and IFN-I promote B cell activation and proliferation (*21, 22*), fostering autoantibody production and IC formation (*23*). NETs also serve as scaffolds for complement activation, amplifying tissue inflammation and damage in lupus (*24, 25*). Thus, the deposition of ICs and C3 in the skin (*26*) or kidneys (*27*) drives local inflammation and tissue damage. Importantly, accumulating evidence indicates that histones directly induce NETosis (*28*) and that NETotic neutrophils trigger secondary NETosis in nearby naive neutrophils (*29*). This self-propagating feedback loop is supported by *in vivo* findings demonstrating a delayed, secondary wave of NET formation in the liver, driven by residual NET components from an initial wave days prior (*30*). This self-sustaining nature underscores the central role of NETosis as both an effector and a driver of systemic autoimmune cascades in SLE.

Building on our published models of UVB-induced acute skin inflammation with neutrophil infiltration and NET formation (*13, 14*), the current study investigates the role of NETosis in acute cutaneous and systemic lupus inflammation using asymptomatic, young female lupus-prone mice―a model that mimics key features of sunlight-triggered lupus flares in patients (*6*). Mechanistically, nuclear envelope rupture is a critical structural step required for NET formation via PKCα-dependent nuclear lamina phosphorylation and disassembly (*13*). Here, we investigated whether inhibiting NETosis by targeting nuclear envelope rupture through *Pkcα* deletion could attenuate skin and kidney inflammation in UVB-irradiated lupus-prone mice. Because only a fraction of neutrophils undergoes NETosis―both *in vitro* (*31, 32*) and in UVB-irradiated skin *in vivo* (*14*)―we employed spatiotemporal photoconversion tracking to monitor the systemic fate of skin-infiltrating neutrophils, including their migration to the kidneys. Critically, it remains unknown whether UVB-induced NETosis and inter-organ neutrophil trafficking act independently or coordinately to link cutaneous photosensitivity to systemic organ injury―a question we address here in the context of acute disease, distinct from the chronic NETosis-driven pathology previously described.

A recent study showed that high-dose UVB induces neutrophil transmigration from the skin to the kidneys, causing transient proteinuria (*33*), however, the effector mechanisms driving this pathology remain undefined. Specifically, the role of NETosis―including its upstream licensing signals, renal propagation, and pathogenic effector cargo that mediates renal injury―remains undefined. To address these gaps, our findings provide the following mechanistic advances. First, we demonstrate that NETosis directly drives local and systemic inflammation. Second, PKCα-dependent nuclear envelope rupture licenses NETosis. Third, NETs serve as scaffolds for effector cytokines (IFNα, IFNγ, IL-17A) and complement C3 in both the skin and kidneys. Fourth, we identify a role for CXCR4 in governing skin-to-kidney neutrophil dissemination, as demonstrated by genetic Cxcr4 deficiency. Collectively, these findings support a two-step model in which local NETosis is followed by the migration of surviving, CXCR4-expressing neutrophils, leading to secondary NETosis in the kidneys. Collectively, these findings position NETs as central effector drivers and unveil new therapeutic strategies for photosensitive SLE.

## RESULTS

### UVB triggered concurrent skin and kidney inflammation with neutrophil infiltration and NET release

Exposure of asymptomatic, young female lupus-prone mice to UVB (Fig 1A) elicited skin inflammation, characterized by increased epidermal and dermal thickness and inflammatory cell infiltration (quantified as nucleated cell count) compared to sham controls (Fig 1B-E). Remarkably, local skin UVB irradiation triggered a systemic response, causing acute renal injury with significant proteinuria and glomerular hypercellularity compared to sham controls (Fig 1F-H). UVB-induced skin inflammation was characterized by robust neutrophil infiltration (Fig 1I) and NET formation (Fig 1J,K) in MRL*/lpr* mice. Notably, these skin-infiltrating neutrophils and their NETs were highly enriched for IFNα, IFNγ, IL-17A, and C3 (Fig 1L-O), establishing NETs as critical reservoirs of proinflammatory effectors in UVB-challenged skin of lupus-prone mice. Crucially, skin inflammation magnitude correlated strongly with systemic disease severity. Specifically, cutaneous inflammatory cell infiltration (nucleated cell count) significantly correlated with proteinuria severity (Fig 1P), a systemic impact further highlighted by the correlation between skin neutrophil infiltration and proteinuria (Fig 1Q). Collectively, these findings suggest that skin-infiltrating immune cells, particularly neutrophils, are not merely local responders but critical pathogenic drivers, potentially mediating an inflammatory cascade from the skin to the kidneys.

**Figure 1.**
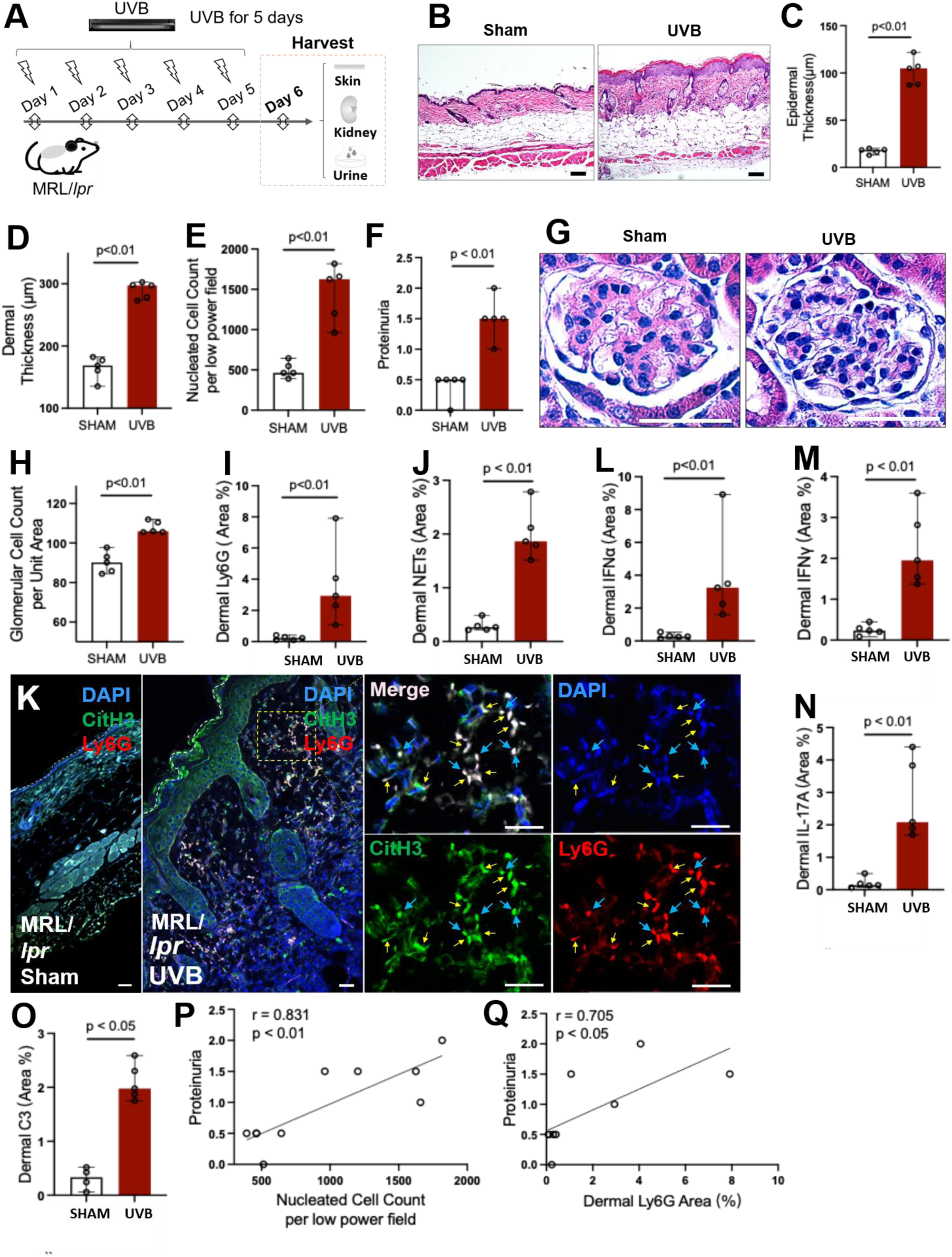
UVB irradiation triggered acute skin and kidney inflammation, neutrophil infiltration and NET formation in young female lupus-prone mice. **A.** Asymptomatic young female MRL/*lpr* mice were irradiated by UVB (150 mJ/cm^2^/day) for 5 days, with urine, skin and kidney samples collected on day 6. **B**-**E**. Representative H&E images (B) and summary data showing increased epidermal (C), dermal (D) skin thickness, and inflammatory cell infiltration (E, nucleated cell count) in the skin of MRL/*lpr* mice without (sham) or with UVB irradiation. **F-H**. Summary of proteinuria scores (F) and glomerular cellularity (H) with representative glomerular H&E images (G) in asymptomatic young female MRL/*lpr* mice without or with UVB irradiation. **I-O**. Summary analysis of skin neutrophils (I), citH3^+^ NETs (J,K), and NET-associated IFNα, IFNγ, IL-17A, and C3 (L-O) in the skin of MRL/*lpr* mice without or with UVB-irradiation. In panels I-O, neutrophils, NETs and NET-associated IFNα, IFNγ, IL-17A, and C3 in kidney sections were stained with PE-labeled anti-ly6G, FITC-labeled antibodies against citH3, IFNα, IFNγ, IL-17A, C3, and DAPI for DNA. Yellow arrows indicate neutrophil NETs (K). **P**-**Q**. Pearson correlations between dermal nucleated cell count (P) or neutrophil infiltration (Q) and proteinuria scores. In panel K, yellow arrows indicate NETs, and light blue arrows indicate non-NETting neutrophils. Scale bars, 100 µm (B,K), and 50 µm (G). Results represent 5 biological replicates per group. Panels C,D-F,H-J,L-O display means ± SD, P<0.05 or <0.01 were determined by Student’s t test for two-group comparisons.

Although previous studies implicate neutrophils in systemic injury following skin thermal burns (*34, 35*) or UVB exposure (*33*), the molecular mechanisms linking cutaneous UVB damage to distant organ damage remain unclear. Given that neutrophil NETosis contributes to both UVB-induced skin inflammation (*13, 14*) and lupus nephritis (*17, 18*), we hypothesized that UVB-triggered NETosis bridges local cutaneous and distant renal inflammation in lupus-prone mice. We therefore investigated the specific role of neutrophil NETosis in driving inflammation across both organs.

### Neutrophil NETosis drives acute kidney inflammation in lupus-prone mice following UVB irradiation

Beyond cutaneous responses (Fig 1I-O), glomerular neutrophil infiltration correlated with glomerular hypercellularity in MRL/lpr mice without or with UVB irradiation (Suppl Fig 1A), suggesting that infiltrating neutrophils contribute to glomerular hypercellularity and renal inflammation. In addition, UVB-irradiated MRL/*lpr* mice exhibited significant glomerular neutrophil infiltration and robust accumulation of NETs (citrullinated H3, citH3) compared to sham controls (Fig 2A-C). Crucially, glomerular NETs co-localized with elevated levels of NET-associated IFNα, IFNγ, IL-17A, and C3 (Fig 2D-H). This glomerular accumulation of neutrophils, NETs, NET-associated cytokines/C3 strongly correlated with the severity of proteinuria in MRL/*lpr* mice (Fig 2I-K, Suppl Fig 1B-D), underscoring its pathological relevance. These results directly link glomerular neutrophil NETs, their associated cytokines, and C3 to acute renal inflammation and injury following UVB irradiation.

**Figure 2.**
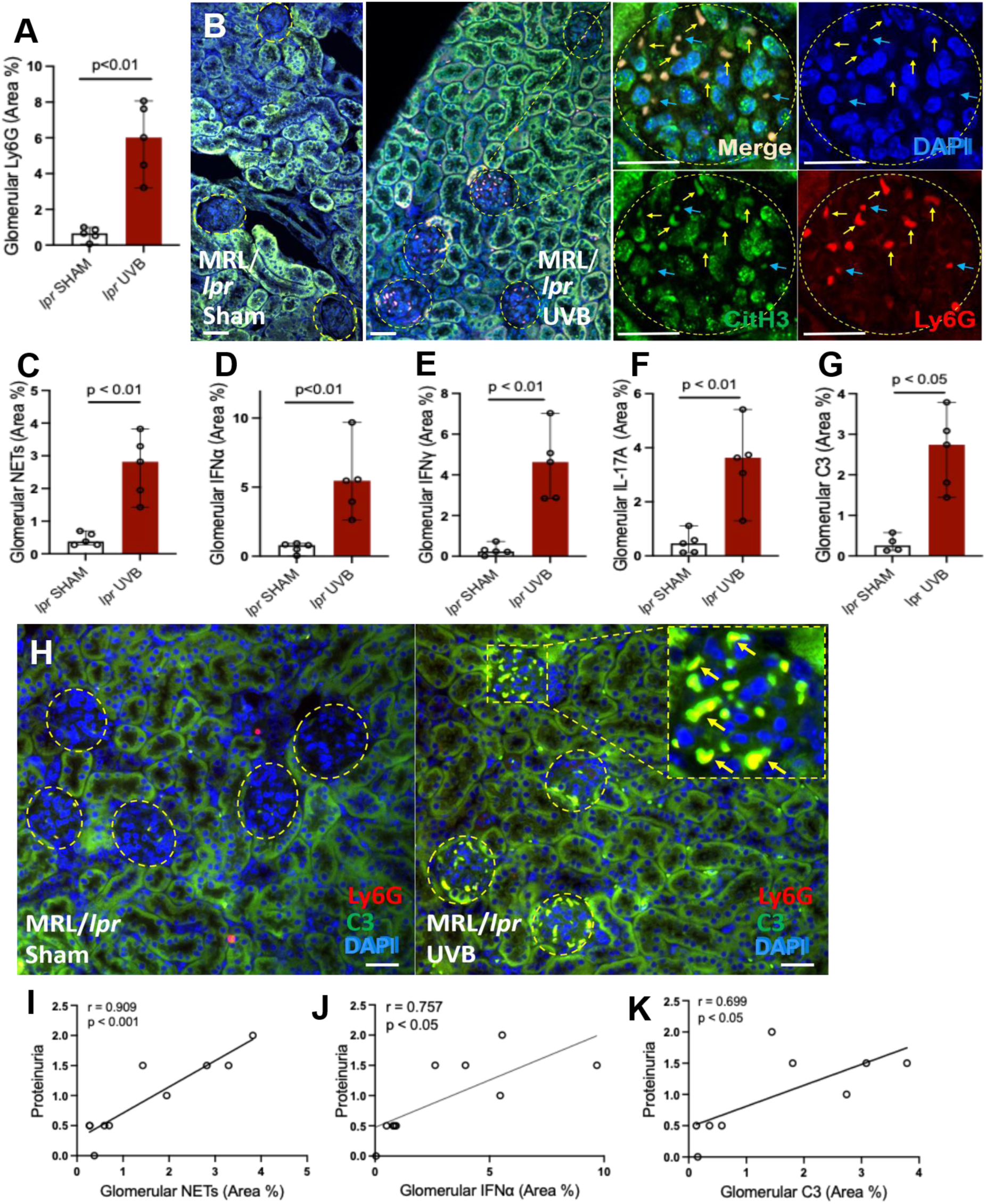
Neutrophil infiltration and NET formation drives acute kidney inflammation in asymptomatic young female lupus-prone mice with UVB irradiation. **A**-**H.** Summary analysis and representative images of neutrophils (A), citH3^+^ NETs (B-C), and NET-associated IFNα, IFNγ, IL-17A, and C3 (D-H), in the kidneys of MRL/*lpr* mice without or with UVB irradiation. Sections were stained with PE-labeled anti-ly6G, FITC-labeled antibodies against citH3, IFNα, IFNγ, IL-17A, C3, and DAPI for DNA. In panels B and H, yellow arrows indicate NETs, and light blue arrows indicate non-NETting neutrophils. Scale bars, 50 µm. **I-K**. Pearson correlations between renal NETs, NET-associated IFNα and C3 deposition in kidneys and proteinuria scores in combined groups of mice. Results represent 5 biological replicates per group. Panels A,C-G display mean ± SD, P<0.05 or P<0.01 were determined by Student’s t test.

Thus, our findings highlight a key role for glomerular-infiltrating neutrophils and their NETs in driving multi-organ inflammation in the skin and kidneys following localized UVB irradiation. Notably, by releasing NETs and associated inflammatory mediators, neutrophils act as critical drivers of coordinated pathogenic responses in both the skin and kidneys of asymptomatic lupus-prone mice. The coordinated responses across both organs (Figs 1-2, Suppl Fig 1) suggest a direct mechanistic link bridging cutaneous and kidney inflammation.

### Targeting nuclear envelope rupture via PKCα genetic deletion attenuated NETosis and UVB-induced lupus skin inflammation

NETosis is an important driver of lupus pathogenesis (*36, 37*), yet the upstream mechanism controlling *in vivo* NET release remains elusive. Since nuclear chromatin externalization and subsequent extracellular release for NET formation require nuclear envelope rupture, our recent cell biology study investigated this structural bottleneck―using both genetic and pharmacological inhibition approaches ―and identified that nuclear envelope rupture is driven by PKCα-mediated nuclear lamin B phosphorylation and disassembly (*13*). The PKCα-Lamin B axis therefore represents a fundamental mechanical requirement for NET formation.

In the current study, stimulating primary neutrophils from lupus-prone mice with either UVB irradiation or platelet-activating factor (PAF)―which is released from UVB-irradiated keratinocytes (*38*)―notably induced PKCα expression. Consistently, combined treatment (PAF + UVB) resulted in a synergistic PKCα upregulation (Suppl Fig 2A), indicating that UVB drives PKCα induction through both direct and indirect pathways. This aligns with previous work showing that PAF activates PKCα in neutrophils (*39*). Therefore, these findings reveal that the pathological environment created by UVB irradiation is uniquely tailored to promote PKCα upregulation in neutrophils. Guided by these findings and our previously defined mechanism of NETosis (*13*), we generated lupus-prone mice with a genetic deletion of *Pkcα* (MRL/*lpr;Pkcα^-/-^*) to investigate how controlling NETosis impacts UVB-induced lupus skin and kidney inflammation.

Following UVB irradiation, MRL/*lpr;Pkcα^-/-^* mice (Fig 3A) exhibited markedly reduced skin inflammation compared to MRL/lpr mice, with decreased epidermal and dermal skin thickness, lower nucleated cell counts, and diminished neutrophil infiltration (Fig 3B-D). Notably, *Pkcα*-deficient lupus-prone mice exhibited significantly reduced NETosis (Fig 3E), corresponding to a lower accumulation of NET-associated IFNα, IFNγ, IL-17A, and C3 in the skin (Fig 3F-K). These findings provide *in vivo* evidence that preserving nuclear envelope integrity via PKCα genetic deletion effectively suppresses NETosis and mitigates cutaneous lupus inflammation. By confirming the critical role of neutrophil NETs and their associated proinflammatory cargo in driving lupus cutaneous pathology, this study highlights nuclear envelope stabilization as a novel therapeutic strategy for lupus.

**Figure 3.**
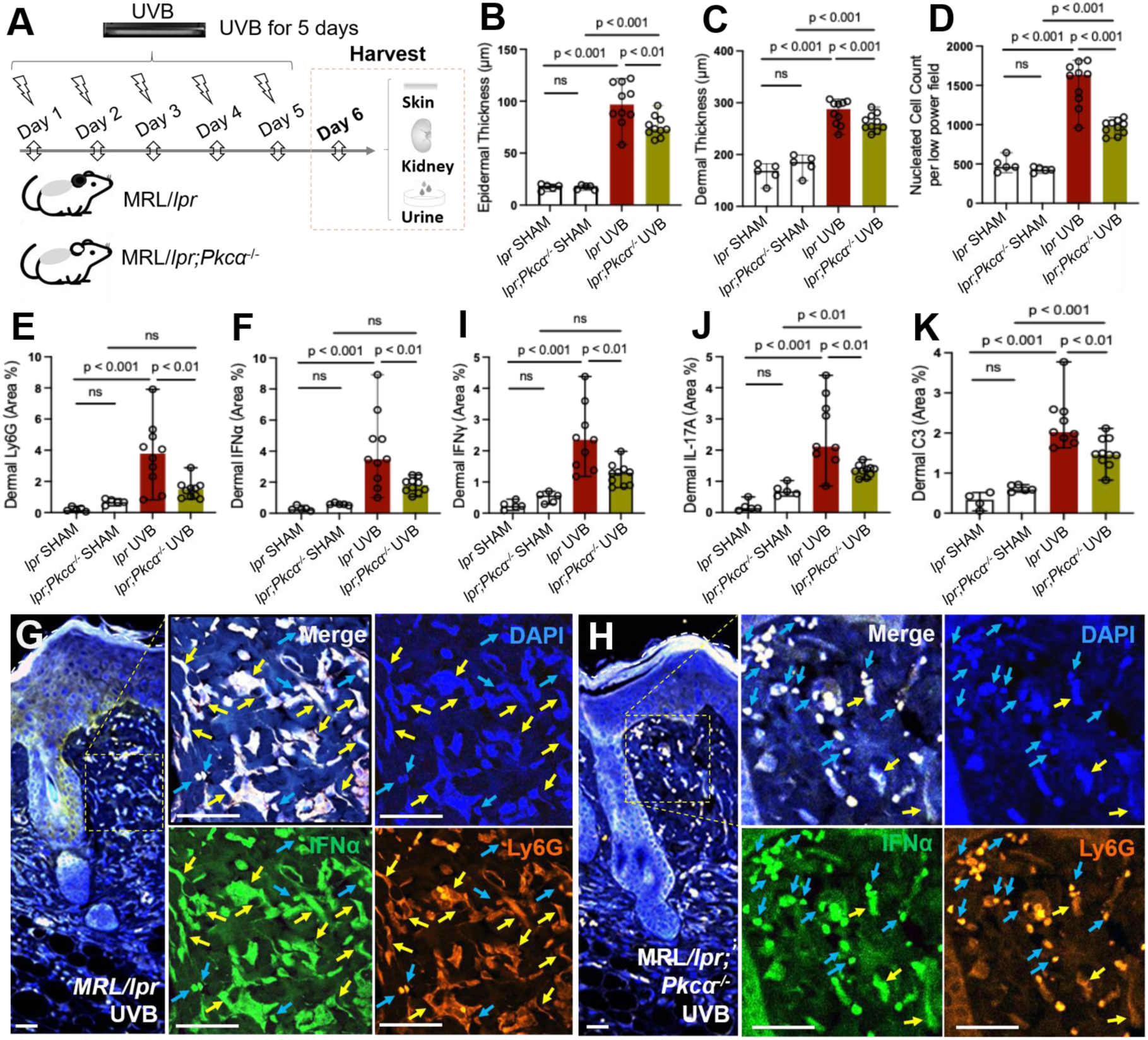
PKCα-dependent nuclear envelope rupture licensed NETosis and amplified UVB-induced skin inflammation in lupus-prone mice. **A**. Asymptomatic female MRL/*lpr* and MRL/*lpr;Pkcα*^-/-^ mice received UVB irradiation (150 mJ/cm^2^/day, 5 days), then urine, skin and kidney samples were harvested on day 6 following cardiac perfusion. **B-D**. Histological quantification of skin thickness of epidermis (B), dermis (C), and proinflammatory cell infiltration (D, nucleated cells) in MRL/*lpr* vs MRL/*lpr;Pkcα*^-/-^ mice without or with UVB irradiation. **E-K**. Analyses of skin neutrophils (E), and NET-associated IFNα (F-H), IFNγ, IL-17A, and C3 (I-K). Skin sections were stained with PE-labeled anti-ly6G, FITC-labeled antibodies against IFNα, IFNγ, IL-17A, or C3, and DAPI for DNA. In panels G and H, yellow arrows indicate NETs, and light blue arrows indicate non-NETting neutrophils. Scale bars, 100 µm. Data represent 5 sham or 10 UVB-irradiated biological replicates. Panels B-F and I-K display mean ± SD; P<0.01 or P<0.001 by one-way ANOVA with SNK post-hoc test.

### Inhibiting PKCα-mediated nuclear envelope rupture suppressed NETosis and ameliorated UVB-induced acute kidney inflammation

Beyond cutaneous effects, our findings demonstrate the profound systemic therapeutic potential of modulating nuclear envelope integrity. Genetic deletion of *Pkcα* significantly inhibited NET formation in glomeruli and decreased UVB-induced acute kidney inflammation and proteinuria in lupus-prone mice (Fig 4A-D). This intervention also led to marked histological improvements, including reduced glomerular hypercellularity (Fig 4B-C) and attenuated neutrophil infiltration (Fig 4D). Critically, we observed a profound reduction in glomerular deposition of neutrophil NETs and NET-associated proinflammatory mediators―including IFNα, IFNγ, IL-17A, and C3―in the kidneys of UVB-irradiated MRL/*lpr;Pkcα^-/-^* mice compared to MRL/*lpr* controls (Fig 4E-I). These data provide genetic evidence that the glomerular deposition of NETs and the associated proinflammatory cargo is a major driver of kidney injury and proteinuria (Fig 2I-K), establishing the targeted regulation of NETosis as a viable therapeutic avenue for photosensitive lupus. Furthermore, our study demonstrates a novel mechanistic paradigm by positioning nuclear envelope integrity as a potent therapeutic target for NETosis and related pathologies, including lupus.

**Figure 4.**
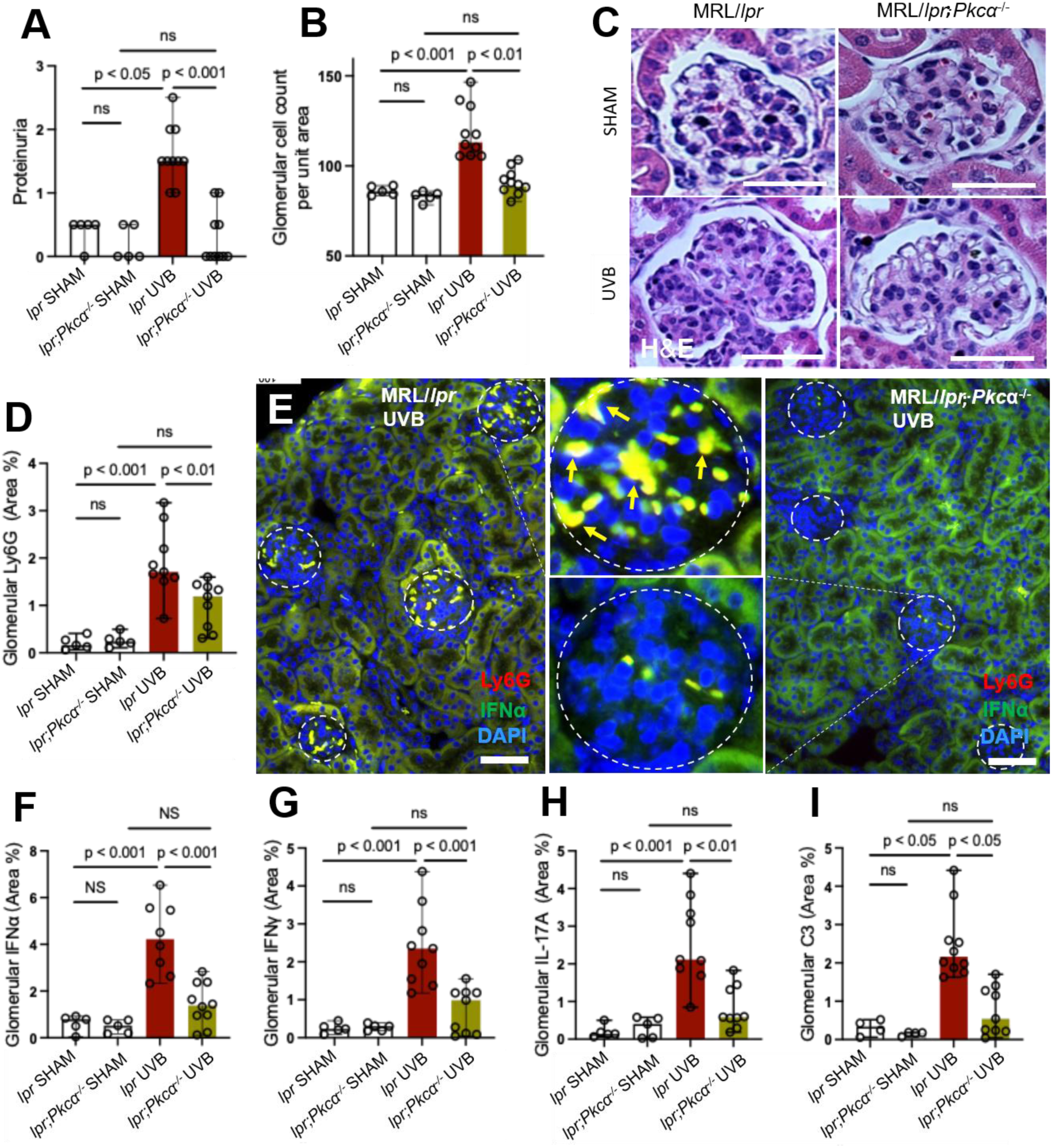
Inhibition of PKCα-dependent NETosis attenuated UVB-induced acute kidney inflammation and proteinuria in lupus-prone mice. **A**-**C**. Quantification of proteinuria scores (A) or glomerular cellularity (B), with representative glomerular images (C) for MRL/*lpr* vs MRL/*lpr;Pkcα*^-/-^ mice without or with UVB irradiation. **D**-**I**. Analysis of glomerular neutrophils (D), and NET-associated IFNα (E,F), IFNγ, IL-17A, C3 (G-I) in MRL/*lpr* vs MRL/*lpr;Pkcα*^-/-^ mice without or with UVB irradiation. Kidney sections were stained with PE-labeled anti-ly6G, FITC-labeled antibodies against IFNα, IFNγ, IL-17A, or C3, and DAPI for DNA. Yellow arrows indicate NETs (E). Scale bars, 50 µm. Data represent 5 sham or 10 UVB-irradiated biological replicates. Panels A,B,D,F-I display mean ± SD; P<0.05, P<0.01, and P<0.001 by one-way ANOVA with SNK post-hoc test.

While our data confirm that NETotic neutrophils are essential effectors of kidney injury, prior studies from our group and others demonstrate that only a fraction of neutrophils undergo NETosis *in vitro* (*31, 32*) and in UVB-irradiated skin *in vivo* (*14*). This divergence introduces an intriguing possibility that surviving neutrophils might act as a migratory cellular bridge, facilitating systemic communication between the inflamed skin and distant renal pathology following UVB irradiation (*33*). To test this hypothesis, we sought to define the mechanistic basis of neutrophil-mediated skin-kidney crosstalk, investigating how these cells coordinate systemic inflammation through neutrophil reverse transmigration and the delivery of NETs, along with their effector proinflammatory cytokines and complement C3.

### UVB irradiation induced a migratory, proinflammatory neutrophil phenotype

To define the mechanisms driving skin-kidney crosstalk, we identify CXCR4-expressing neutrophils as a critical cellular link mediating inter-organ communication during UVB-induced acute systemic lupus exacerbation. Strikingly, compared with sham controls, UVB irradiation triggered a marked systemic expansion of CXCR4-expressing neutrophils in both the skin and kidneys, regardless of their NETotic status, with CXCR4 also detected within released NETs (Fig 5A-D). Crucially, UVB-irradiation significantly increased the pool of circulating CXCR4-expressing neutrophils in the peripheral blood (Fig 5E,F), consistent with the systemic mobilization and dissemination of this population. These observations support a model in which CXCR4-expressing neutrophils serve as a pathogenic bridge between inflamed skin and distant kidneys, promoting the inter-organ propagation of inflammation in photosensitive lupus.

**Figure 5.**
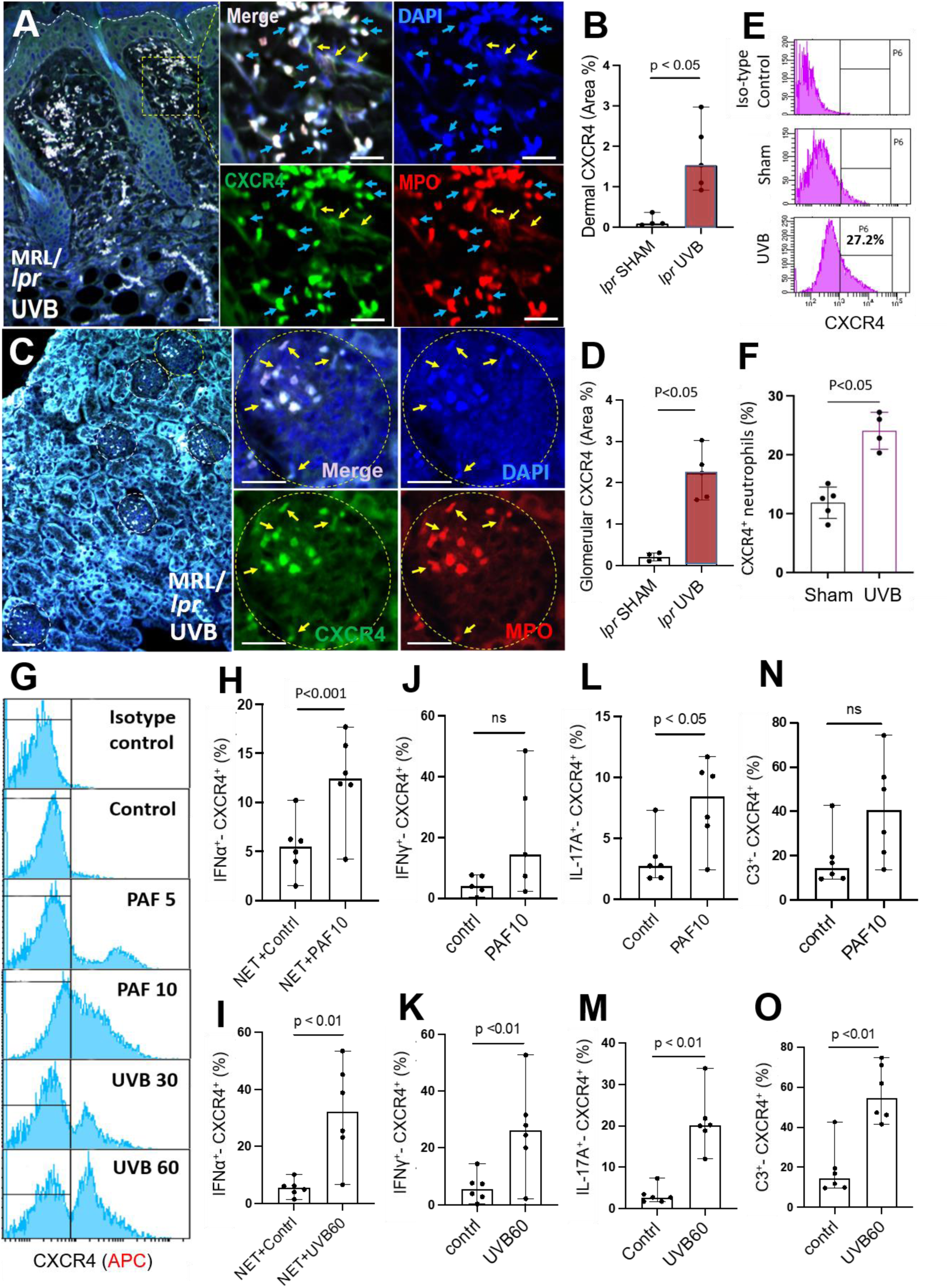
UVB-induced a migratory, proinflammatory CXCR4-expressing neutrophil phenotype. **A-D**. Immunofluorescence images and summary analysis of co-expression of MPO and CXCR4 and the deposition of neutrophil NETs in the inflamed skin and kidneys of UVB-irradiated MRL/*lpr* mice. Skin and kidney sections were stained with Alexa 647-labeled MPO, FITC-labeled antibodies against CXCR4, and DAPI for DNA. In panels A and C, yellow arrows indicate NETs, and light blue arrows indicate non-NETting neutrophils. Scale bars, 100 µm (skin) or 50 µm (kidneys). **E**,**F**. Flow cytometry histograms of CXCR4 expression in peripheral blood neutrophils from MRL/*lpr* mice without or with UVB irradiation (E,F). Peripheral blood neutrophils were stained with PE-labeled anti-mouse ly6G and APC-labeled rat anti-mouse CXCR4 or APC-labeled rat IgG2b isotype control. **G**-**O**. Representative histogram and summary analyses of flow cytometry data for primary neutrophils from MRL/*lpr* mice stimulated *in vitro* with PAF (10 µM PAF) or UVB (60 mJ/cm^2^) and incubated for 20h. Panels show neutrophil expression of CXCR4 (G), and cytokines IFNα (H,I), IFNγ (J,K), IL-17A (L,M), and C3 (N,O). Cells were stained with PE-labeled anti-mouse ly6G, APC-labeled anti-mouse CXCR4, and FITC-labeled anti-mouse IFNα, IFNγ, IF-17A, or C3. All neutrophils in panels E-O were gated on ly6G^+^ cells. Data represent 4-6 biological replicates. Panels B,D,F,H-O display means ± SD, P<0.05, P<0.01, P<0.001 determined by Student’s t test for two-group comparisons.

To delineate the underlying mechanisms, we treated primary neutrophils isolated from MRL/*lpr* mice with UVB irradiation or PAF. Both stimuli robustly upregulated CXCR4 expression in neutrophils in a dose-dependent manner (Fig 5G). This phenotypic upregulation was accompanied by elevated expression of key inflammatory mediators, including IFNα (in the presence of DNA-NETs, as previously characterized (*13, 14*)), IFNγ, IL-17A, and C3 (Suppl Fig 3A-D). Notably, a subset of these primed neutrophils co-expressed CXCR4 concurrently with these proinflammatory mediators (Fig 5H-O, Suppl Fig 3E-H). These findings demonstrate that UVB exposure―acting through both direct cellular stress and indirect PAF paracrine signaling―reprograms skin-infiltrating neutrophils, empowering them with a dual migratory-inflammatory capability.

The co-expression of CXCR4 and cytokines/C3 enables specific neutrophil subsets to migrate from inflamed skin to kidneys. Recognizing only a fraction of neutrophils undergoes NETosis―both *in vitro* (*31, 32*) and *in vivo* in UVB-irradiated skin (*14*)―we propose that UVB-primed neutrophils adopt functionally distinct yet overlapping states that enable local inflammation and systemic dissemination. One subset of neutrophils undergoes terminal NETosis (*13*) to drive local skin inflammation via effector NET-associated cytokines/C3. In parallel, the surviving, CXCR4-expressing neutrophils retain migratory capability, undergo reverse transmigration (RTM) to re-enter the circulation (*40, 41*), and are recruited to the kidneys. This recruitment is driven by elevated renal CXCL12 levels, which occur either following UVB irradiation (*33*) or intrinsically in various lupus-prone strains, including MRL/*lpr* mice (*42*), and in patients with lupus nephritis (*43*). Importantly, these phenotypic states are not mutually exclusive, reflecting a continuum of activation rather than a strictly binary fate decision. This overlapping phenotype underscores our proposed two-step model, wherein UVB irradiation induces both local NETosis and the generation of a migratory pool of primed, CXCR4-expressing neutrophils poised for systemic dissemination. Upon sequestration within the glomerular microenvironment, these primed neutrophils undergo secondary NETosis, releasing NET-associated cytokines/C3 that were likely synthesized before their migration from the UVB-irradiated skin, thereby transducing a cutaneous insult into renal inflammation.

### Spatiotemporal tracking identified a critical role for CXCR4 in skin-kidney neutrophil transmigration

To causally link skin UVB irradiation to renal injury, we employed precise spatiotemporal photoconversion tracking using *kikGR;Cxcr4*^+/-^ mice, which was generated by crossing Cxcr4-deficient mice with photoconvertible reporter *kikGR*-expressing mice (Fig 6A), enabling "tagging" skin neutrophils with violet light to monitor their migration across other organs. These mice enable precise spatiotemporal *in vivo* tracking of neutrophil migration (*44*), directly revealing the impact of CXCR4 deficiency on cellular trafficking. Sequential UVB and violet light irradiation increased photoconverted kikR^+^ neutrophils in the circulation as detected by flow cytometry (Fig 6B). Critically, Cxcr4 deficiency significantly reduced circulating kikR^+^ neutrophils in *kikGR;Cxcr4*^+/-^ mice compared to *kikGR;Cxcr4^flox^*controls (Fig 6B), demonstrating the essential role of CXCR4 in neutrophil RTM from UVB-irradiated skin into circulation.

**Figure 6.**
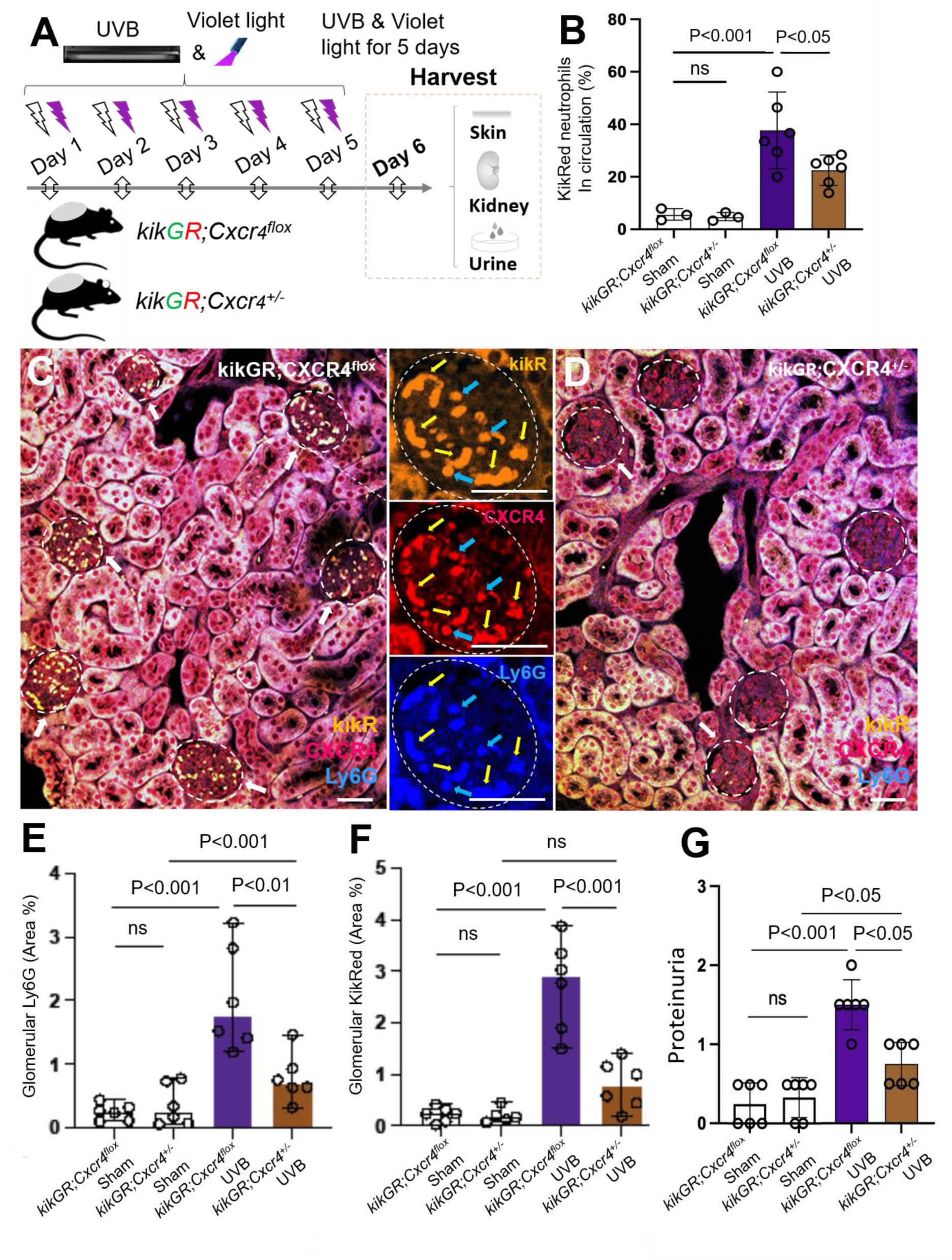
CXCR4 mediates neutrophil systemic dissemination from the skin to the kidneys. **A.** Young *kikGR;Cxcr4*^+/-^ and *kikGR;Cxcr4^flox^*control mice expressing a photo-convertible reporter were irradiated by UVB (150 mJ/cm^2^/day) and violet light for 5 days, then urine, skin and kidney samples were harvested on day 6. **B**. Flow cytometry analysis of peripheral blood kikR^+^ neutrophils from *kikGR;Cxcr4^flox^* and *kikGR;Cxcr4*^+/-^ mice without or with UVB and violet light irradiation. Analysis was performed to assess systemic circulation, gating on Ly6G^+^ cells (PE-labeled anti-mouse ly6G) that had photoconverted from kikG^+^ to kikR^+^. **C**-**F**. Immunofluorescence and summary analysis of photoconverted kikR^+^ neutrophil infiltration into kidneys. Kidney sections were stained with Pacific Blue labeled anti-mouse ly6G, and Alexa 647-labeled CXCR4. White arrows indicate glomeruli with deposition of photoconverted kikR^+^ neutrophils, while blue and yellow arrows denote deposition of kikR^+^ /CXCR4^+^ neutrophils and NET-like structures within glomeruli, respectively. Scale bars, 50 µm. **G**. Quantification of proteinuria scores in control *kikGR;Cxcr4^flox^* and *kikGR;Cxcr4*^+/-^ mice without or with UVB and violet light irradiation. Data represent 6 biological replicates in each group (3 male and 3 female mice). Panels E-G display mean ± SD, P<0.05, P<0.01, and P<0.001 determined by one-way ANOVA with SNK post-hoc test.

Histological analysis revealed significant accumulation of ly6G^+^ and kikR^+^ neutrophils and NET-like structures in glomeruli of UVB-irradiated control *kikGR;Cxcr4^flox^* mice following violet light exposure (Fig 6C,E,F). This accumulation (ly6G^+^ and kikR^+^ neutrophils) was significantly reduced in *Cxcr4*-deficient *kikGR;Cxcr4*^+/-^ mice (Fig 6D-F). Consequently, decreased pathogenic neutrophil dissemination resulted in significantly reduced proteinuria in UVB-irradiated *kikGR;Cxcr4*^+/-^ mice compared to controls (Fig 6G). These data provide direct evidence that CXCR4 is a causal mediator guiding pathogenic neutrophil dissemination from inflamed skin to kidneys, functionally demonstrating that the CXCR4-mediated skin-kidney neutrophil dissemination drives acute kidney inflammation and subsequent renal injury. These findings provide mechanistic evidence linking UVB-induced acute skin inflammation to systemic lupus exacerbation. Specifically, we demonstrate that CXCR4-mediated skin-kidney neutrophil trafficking is essential for the systemic dissemination of inflammatory signals from localized skin lesions to the kidneys, where they undergo secondary NETosis to drive renal injury.

### Pharmacological inhibition of CXCR4 attenuated renal inflammation following UVB-irradiation

Building on genetic evidence, we further evaluated the translational potential of pharmacological CXCR4 inhibition using the selective antagonist IT1t in UVB-irradiated MRL/*lpr* lupus-prone mice (Fig 7A). Pharmacological CXCR4 inhibition significantly suppressed the acute renal sequelae of UVB irradiation, yielding marked reductions in proteinuria, glomerular hypercellularity, and glomerular infiltration of Ly6G^+^ and CXCR4^+^ neutrophils (Fig 7B-F). Moreover, *in vitro* transwell migration assays confirmed that IT1t significantly reduced neutrophil migratory capacity (Fig 7G), validating its role in blocking neutrophil recruitment to inflamed tissues.

**Figure 7.**
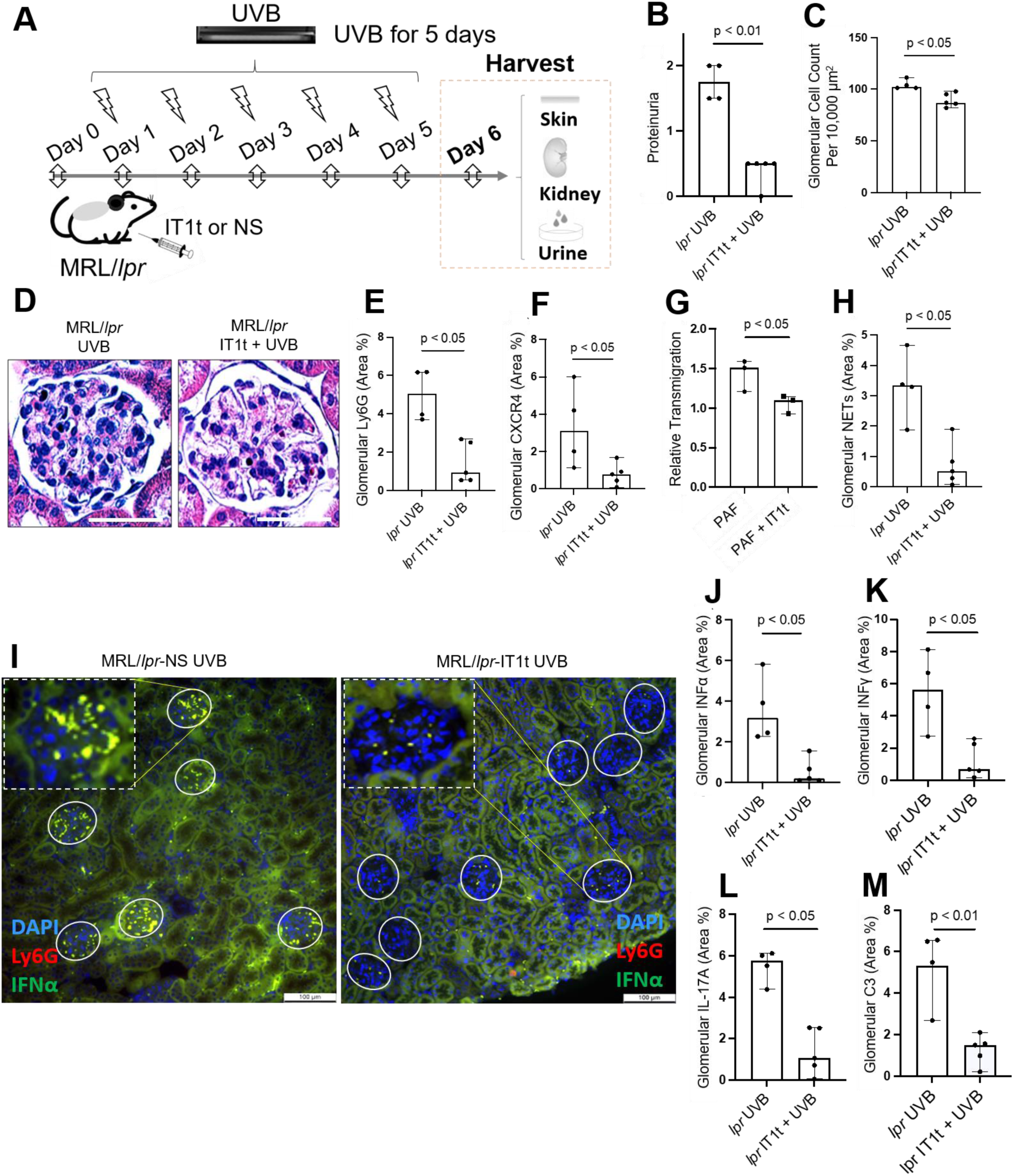
Pharmacological CXCR4 blockade interrupts the NETosis-driven skin-kidney inflammatory axis. **A**. Asymptomatic female MRL/*lpr* mice irradiated by UVB (150 mJ/cm^2^/day, 5 days), with intraperitoneal administration of CXCR4 antagonist IT1t (10 mg/kg/day, 6 days) began one day prior to the first session of UVB irradiation Then, urine, skin and kidney samples were harvested on day 6 post cardiac perfusion. **B**-**F**. Quantification of proteinuria scores (B), glomerular cellularity (C,D), renal Ly6G-neutrophils (E) and CXCR4-expressing neutrophils (F) in UVB-irradiated MRL/*lpr* mice without (control) or with IT1t administration. **G**. Relative migration analysis of mouse neutrophils with CXCR4 expression, induced by 10 µM PAF stimulation for 20h, was conducted using a Boyden chamber. Cells were pretreated without or with IT1t, and then stimulated with CXCL12 at 300 ng/ml. **H**-**M**. Analyses of glomerular deposition of neutrophil NETs, and NET-associated IFNα, IFNγ, IL-17A, or C3 in glomeruli of UVB-irradiated MRL/*lpr* mice without (control) or with IT1t pre-administration. Sections (E,F,H-M) were immunostained with PE-labeled anti-ly6G, FITC-labeled antibodies against CXCR4, citH3, IFNα, IFNγ, IL-17A, or C3, and DAPI for DNA. Scale bars, 50 µm (D), and 100 µm (I). Data represent 3-5 biological replicates. Panels B,C,E-H,J-M display means ± SD, P<0.05, P<0.01 by Student’s t test for two-group comparisons.

**Figure 8.**
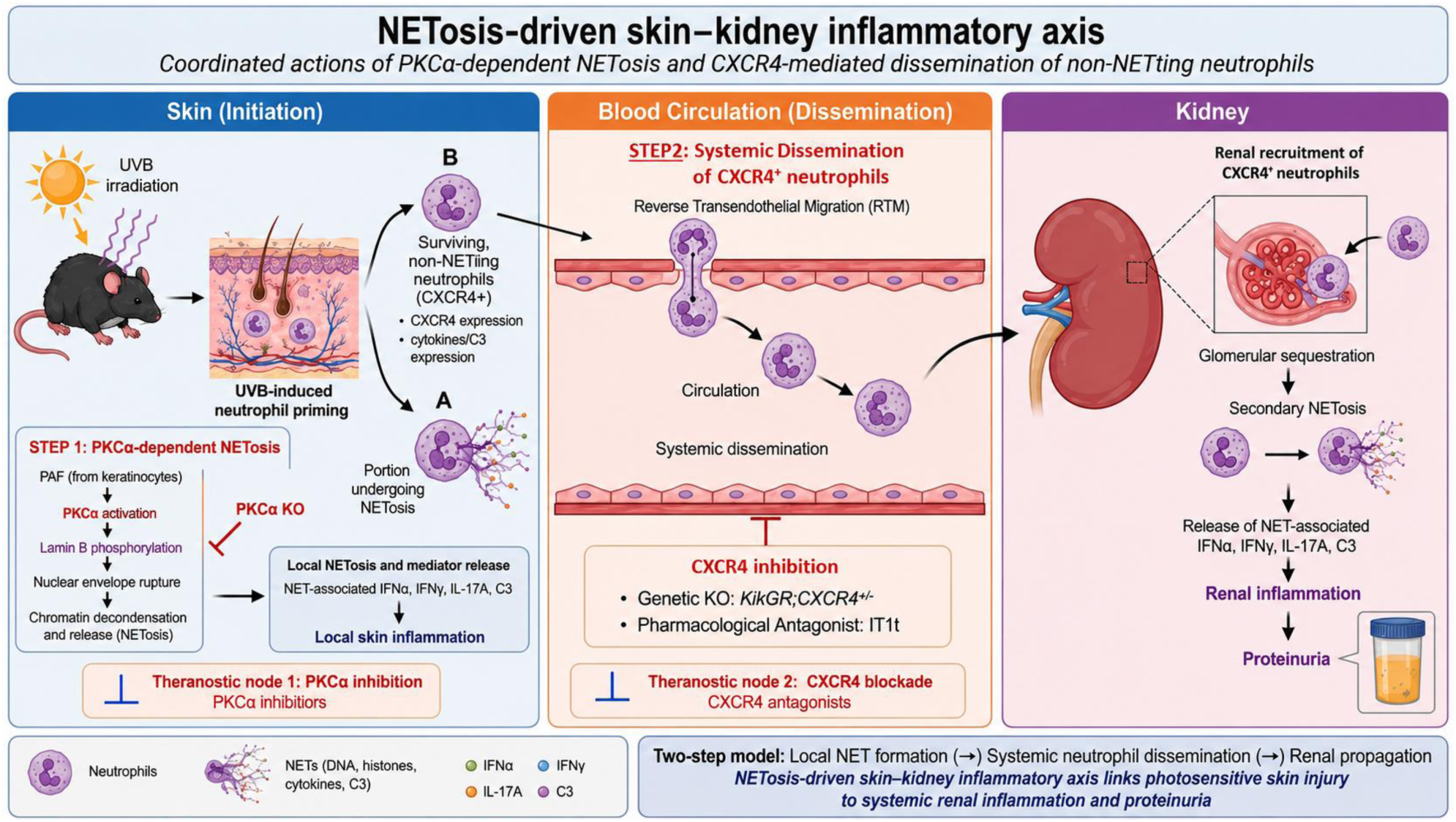
Schematic model of a NETosis-driven skin–kidney inflammatory axis in photosensitive lupus. UVB irradiation induces neutrophil infiltration and priming within the inflamed skin, resulting in two distinct neutrophil fates. ***i)*** A subset of skin-infiltrating neutrophils undergoes PKCα-dependent NETosis (**Step 1, Initiation)**, in which platelet-activating factor (PAF, released from UVB-irradiated keratinocytes) activates PKCα, leading to lamin B phosphorylation and disassembly, nuclear envelope rupture, and NET formation, and accompanied by the local release of effector NET-associated proinflammatory cytokines (IFNα, IFNγ, IL-17A) and complement C3, thereby promoting cutaneous inflammation. Genetic deletion of PKCα suppresses local inflammatory responses through inhibiting nuclear envelope rupture and NET formation. ***ii)*** A second population of surviving, non-NETting neutrophils acquires a migratory, proinflammatory phenotype characterized by upregulation of CXCR4, cytokines and complement C3. These CXCR4-expressing neutrophils undergo reverse transendothelial migration (**Step 2, Systemic dissemination**), re-enter the circulation, and subsequently home to the kidneys, where they accumulate in glomeruli and undergo secondary NETosis. The release of NET-associated cytokines and complement C3 in the kidney propagates renal inflammation and proteinuria. Together, PKCα-dependent NETosis and CXCR4-mediated dissemination of surviving neutrophils establish a self-amplifying, NETosis-driven skin– kidney inflammatory axis linking photosensitive skin injury to systemic renal inflammation. The model further identifies two complementary therapeutic opportunities for intercepting UVB triggered lupus flares: inhibition of PKCα-dependent NETosis and blockade of CXCR4-mediated neutrophil dissemination.

Mechanistically, systemic pharmacological CXCR4 inhibition significantly attenuated acute kidney inflammation in UVB-irradiated MRL/*lpr* mice, as evidenced by the reductions in proteinuria and glomerular hypercellularity, and blocked neutrophil infiltration into the kidneys. Furthermore, CXCR4 inhibition ameliorated glomerular accumulation of NETs, and NET-associated IFNα, IFNγ, IL-17A, and C3 compared to saline-treated control MRL/*lpr* mice (Fig 7H-M). Therefore, these findings reveal a novel pathogenic link between UVB irradiation and renal damage, highlighting that CXCR4-mediated neutrophil recruitment drives acute kidney inflammation via NET formation and the release of effector NET-associated inflammatory mediators.

In addition to its effects on the kidneys, CXCR4 inhibition with IT1t attenuated dermal (but not epidermal) thickness and reduced nucleated cell counts in UVB-irradiated MRL/lpr mice (Suppl Fig 4A-D). This was driven by a significant decrease in Ly6G neutrophil infiltration, NET formation, and NET-associated IFNα and IL-17A (Suppl Fig 4E,G,H,J). However, CXCR4-expressing neutrophils, NET-associated IFNγ and C3 in the skin did not show significant differences between IT1t-treated mice and saline control mice (Suppl Fig 4F,I,K). This efficacy of CXCR4 inhibition―achieved via repeated systemic administration of IT1t―underscores a critical role for CXCR4 in coordinating systemic inflammatory responses to UVB, aligning with previous reports where CXCR4 inhibition attenuated sunlight-induced skin cancer by preventing mast cell infiltration into the skin (*45*).

Collectively, our findings elucidate a novel pathogenic mechanism driving UVB-induced systemic inflammation in photosensitive lupus. We demonstrate that a NETosis-driven skin–kidney inflammatory axis is established through the coordinated actions of PKCα-dependent NETosis and CXCR4-mediated dissemination. This process is governed by CXCR4-dependent neutrophil trafficking from UVB-irradiated skin to the kidneys, where recruited neutrophils undergo secondary NETosis and release their effector NET-associated cytokines/C3. By establishing this causal, two-step axis, our work identifies PKCα-dependent nuclear envelope rupture and CXCR4-mediated neutrophil dissemination as coordinated, mechanistically distinct therapeutic targets. These specific pathways offer a high-precision strategy to uncouple environmental triggers from end-organ damage and mitigate acute systemic inflammation in photosensitive lupus.

## DISCUSSION

Sunlight overexposure exacerbates lupus by inducing acute skin inflammation and systemic disease, including lupus nephritis (*5–8*). Whereas previous studies established NETosis as a contributor to chronic lupus pathogenesis (*15–18*), here we identify a previously unrecognized, acute NETosis-driven skin–kidney inflammatory axis. Our findings demonstrate that PKCα-dependent NETosis serves as the initiating event, while CXCR4-mediated dissemination of surviving neutrophils propagates inflammation to distant kidneys. Beyond prior reports on skin-kidney neutrophil trafficking (*33*), we demonstrate that NETosis—not merely neutrophil infiltration—is the pathogenic effector, with renal NET deposition correlating with proteinuria and both attenuated upon *Pkcα* deletion. Mechanistically, UVB induces terminal NETosis in a subset of skin-infiltrating neutrophils to drive local inflammation, while surviving CXCR4-expressing neutrophils transmigrate to the glomeruli, where they undergo secondary NETosis and release NET-associated proinflammatory cytokines and C3, which drive renal injury and proteinuria. This two-step pathogenic, self-amplifying inflammatory axis reveals two complementary, non-redundant therapeutic targets for photosensitive lupus.

Using initially asymptomatic, young female lupus-prone mice, we show that infiltrating neutrophils and their NETs deposit proinflammatory cytokines and C3 in both skin and kidneys, driving local inflammation and proteinuria. Although NETosis is a recognized contributor to chronic lupus pathogenesis (*36, 37*), its role in acute, UVB-triggered flares remained unclear. Our *in vivo* evidence identifies NETosis as a key effector of UVB-induced acute inflammation in both skin and kidney. We demonstrate that NETs bearing pro-inflammatory cytokines and C3 deposit in both organs, with their glomerular levels correlating strongly with kidney injury and proteinuria. Unlike soluble cytokines and C3 secreted by neutrophils―which rapidly diffuse and are diluted by circulating blood (*14*)―NETs serve as scaffolds that concentrate these pathogenic effectors locally within the skin and kidneys (*14*), driving tissue injury. Moving beyond previous studies of neutrophil trafficking (*33*), our findings position NETs and NET-associated effectors as pathogenic drivers of a self-amplifying cycle, further exacerbated by NET remnants that trigger secondary NETosis in naive neutrophils *in vitro* (*29*) and *in vivo* (*30*). Thus, we extend the pathogenic roles of NETosis from chronic lupus (*15, 16*) and UVB-induced skin inflammation (*13, 14*) to acute systemic responses to sunlight.

Importantly, this work identifies PKCα-mediated nuclear envelope rupture as a critical, non-redundant licensing step in the NETosis-driven skin–kidney axis. Although peptidyl-arginine deaminase 4 (PAD4)-mediated chromatin decondensation has been considered central to NETosis (*37, 46*), its therapeutic relevance remains controversial. Early studies showed PAD4 inhibition attenuated skin and kidney inflammation in aged lupus-prone mice (*15*), however subsequent genetic and pharmacological investigations failed to protect against lupus skin and systemic disease in similar models (*47*). These discrepancies underscore the limitations of targeting PAD4 and highlight the need to target more fundamental, obligatory mechanisms governing NET formation.

Recent mechanistic studies have identified that nuclear chromatin decondensation and nuclear envelope rupture are essential cellular events required for chromatin release and NET formation (*37, 46*). As the primary physical barrier sequestering nuclear chromatin, the nuclear envelope relies on the structural integrity of the nuclear lamina, composed of lamin A/C and B (*37, 46*). Regarding the breakdown of this barrier, while neutrophil elastase is known to be involved in NET formation (*32*), there is no evidence identifying lamins as direct substrates of neutrophil elastase (*48*). Intriguingly, we have identified that nuclear envelope rupture and NET formation are driven by PKCα-mediated phosphorylation and subsequent disassembly of nuclear lamin B, rather than proteolytic cleavage (*13, 14, 37*).

The current study links this pathway to photosensitivity-induced systemic inflammation. UVB irradiation induces keratinocyte production of PAF (*38*), which activates neutrophil PKCα signaling (*39*) in the literature. Here, stimulation of primary neutrophils from lupus-prone mice with either UVB or PAF significantly increased PKCα expression, while combined UVB and PAF treatment yielded a synergistic increase in PKCα levels. Thus, UVB promotes PKCα induction through both direct and indirect pathways, creating a cutaneous inflammatory microenvironment tailored to license NETosis. In this model, UVB-induced PAF amplifies PKCα signaling to drive lamin B phosphorylation, nuclear envelope rupture, and subsequent neutrophil NET formation within the skin of UVB-irradiated mice. This phosphorylation-dependent mechanism represents a structural prerequisite for NET formation, and provides a mechanistically distinct therapeutic target upstream of nuclear chromatin release. Using MRL/*lpr;Pkcα^-/-^*mice generated by backcrossing the *Pkcα* deficiency onto the MRL/*lpr* background for over ten generations to ensure genetic purity, we provide in vivo evidence that preserving nuclear envelope integrity through *Pkcα* genetic deletion effectively suppressed NETosis and significantly ameliorated UVB-induced skin and kidney inflammation compared to control MRL/*lpr* mice. These findings provide the first in vivo mechanistic evidence supporting the novel concept that nuclear envelope regulation represents a therapeutically tractable strategy for mitigating NETosis-driven pathology in lupus skin-kidney axis. They further highlight NETosis as a potential therapeutic target for acute systemic inflammation in photosensitive lupus.

A pathological skin-kidney connection has long been suggested by clinical observations, including the onset of proteinuria following severe skin thermal burns (*49*), with activated neutrophils implicated through the release of neutrophil gelatinase-associated lipocalin (*35*). This systemic link is further supported by a recent mouse study showing that a single high-dose UVB induces transient proteinuria alongside neutrophil transmigration from inflamed skin to kidneys (*33*). Collectively, these studies implicate neutrophils as key mediators of a conserved skin-kidney axis linking acute skin injury to kidney inflammation (*33, 35*). Extending this paradigm, our study provides direct evidence for an acute NETosis-driven skin-kidney inflammatory axis triggered by UVB irradiation in lupus. The severity of skin inflammation was strongly associated with renal injury and proteinuria, supporting a coordinated systemic process rather than two independent organ pathologies and identifying neutrophils and their NETs as critical mediators of acute skin-kidney crosstalk during UVB-induced lupus exacerbation.

Our data established a causal mechanistic link between UVB-induced skin inflammation and systemic disease dissemination through CXCR4-mediated neutrophil trafficking. These findings extend previous clinical observations that circulating leukocytes from SLE patients exhibit increased CXCR4 expression (*43, 50, 51*), suggesting that dysregulated CXCR4 signaling represents a hallmark of systemic lupus immune activation. UVB primed skin-infiltrating neutrophils to upregulate CXCR4, a key mediator of inter-organ trafficking (*40, 41*), linking our findings to the CXCR4-CXCL12 axis implicated in lupus progression (*42, 43, 50–54*). Consistent with prior *in vitro* and *in vivo* observations (*31, 32, 55*) and our prior work (*14*), only a subset of neutrophils undergoes NETosis at the primary site. Thus, CXCR4 functions as a molecular checkpoint enabling surviving, UVB-primed neutrophils to exit inflamed skin, re-enter circulation, and home to kidneys with CXCL12 expression (*33, 42, 43, 52, 54*). Renal CXCL12 expression is elevated in kidneys of both lupus nephritis patients and various lupus-prone mice, creating a chemotactic environment that promotes recruitment of CXCR4-expressing immune cells (*42, 43, 52, 54*). UVB further enhances this susceptibility by inducing acute renal CXCL12 expression (*33*). Collectively, these findings support a mechanism by which UVB-induced CXCR4 upregulation converts localized skin inflammation into systemic renal injury, underscoring a critical role for skin-infiltrating neutrophils in driving lupus kidney inflammation and injury (*56*). Mechanistically, UVB, directly or through PAF signaling (*57*), primes skin-infiltrating neutrophils to express CXCR4 alongside pathogenic mediators (IFNα, IFNγ, IL-17A, C3), thereby generating a migratory, proinflammatory neutrophil phenotype that bridges skin inflammation and kidney injury.

Once sequestered in the kidneys, these primed neutrophils undergo secondary NETosis, releasing NET-associated cytokines/C3―likely synthesized in UVB-irradiated skin―to drive acute kidney inflammation and proteinuria. Although NETosis typically occurs within 3 hours of stimulation (*13, 14, 37*), delayed NETosis has been observed following UVB (4-6 hours) (*13*) and PMA, LPS, or G-CSF (24-48 hours) stimulation (*58*). Consistent with this temporal flexibility, a delayed second wave of NETosis has recently been demonstrated in vivo several days after LPS challenge, driven by vascular NET remnants from the initial wave (*30*), providing a temporal window for neutrophil RTM and renal homing before terminal NET formation (*30*). Together, these findings support a two-step model in which localized NETosis initiates cutaneous inflammation, whereas surviving CXCR4-expressing neutrophils disseminate to the kidneys and undergo secondary NETosis, releasing NET-associated cytokines/C3 to drive renal inflammation and proteinuria. This coordinated sequence establishes a pathogenic, NETosis-driven skin–kidney inflammatory axis linking UVB-induced photosensitivity to systemic lupus flares.

Using the unique spatiotemporal tracking capability of the *kikGR;Cxcr4*^+/-^ mouse model, we provide *in vivo* evidence that CXCR4 governs the egress of activated neutrophils from UVB-irradiated skin back into the circulation and their subsequent accumulation in renal glomeruli. These findings establish a causal skin-kidney axis in which cutaneous UVB irradiation drives renal neutrophil infiltration, NET formation, and proteinuria. Mechanistically, Cxcr4 deficiency impaired neutrophil RTM, reducing the systemic pool of pathogenic neutrophils available for renal recruitment. Thus, our data identify the CXCR4-CXCL12 axis as controlling neutrophil re-entry into the circulation and re-deposition in the kidneys. Although our study focuses on acute UVB effects, these findings align with broader SLE evidence, including increased CXCR4 expression on immune cells and elevated renal CXCL12 levels in SLE patients and lupus-prone mice (*42, 43, 50, 52, 54*).

By identifying an essential role of CXCR4 in neutrophil RTM, we demonstrate that localized skin inflammation can drive distant organ injury in photosensitive lupus. Genetic Cxcr4 deficiency blocks neutrophil re-entry into circulation, reducing the systemic pool of primed pathogenic neutrophils and establishing a mechanistic pathway linking cutaneous UVB irradiation to renal neutrophil accumulation, NET formation, and proteinuria. To translate this mechanistic insight, we evaluated the selective CXCR4 antagonist IT1t in UVB-irradiated lupus-prone mice using a validated dosing strategy in other lupus models (*59*). CXCR4 inhibition blocked renal neutrophil recruitment and also reduced dermal thickness and neutrophil infiltration without affecting the epidermis, confirming that CXCR4 signaling is essential for immune cell migration into the skin dermis (*45*). While previous studies identified CXCR4 as a negative regulator of TLR7-mediated IFN-I signaling in plasmacytoid dendritic cells (*59*), our work expands this framework by showing that CXCR4 inhibition also blocks pathogenic neutrophil dissemination to the kidneys.

By identifying CXCR4 as the critical mediator of skin-kidney trafficking, our study suggests a therapeutic paradigm shift in photosensitive SLE―from broad immunosuppression toward targeted blockade of CXCR4-mediated neutrophil dissemination. This strategy offers a more precise approach to uncouple localized skin inflammation from systemic disease exacerbation, interrupting progression to internal organ injury. Importantly, CXCR4 can be pharmacologically targeted with existing FDA-approved inhibitors, such as AMD3100/plerixafor, which block pathogenic neutrophil dissemination to the kidneys with the potential to avoid broad immunosuppression. Since UVB-induced proteinuria occurs in both lupus-prone (current study) and non-lupus mice (*33*), this pathway may represent a general mechanism linking photosensitivity to systemic inflammatory injury, with potential relevance to other autoimmune or inflammatory conditions.

Our findings support an integrated, two-step model of disease propagation in photosensitive SLE. In this model, PKCα-dependent NETosis drives local inflammation, while CXCR4-mediated dissemination of surviving neutrophils propagates inflammation to the kidneys, where secondary NETosis sustains renal injury. Beyond previous observation of skin-kidney neutrophil transmigration (*33*), we show that these migrating neutrophils deliver effector NETs―scaffolded with proinflammatory cytokines/C3―as pathogenic mediators of renal inflammation and injury. Together, these coordinated processes establish a self-amplifying skin–kidney inflammatory axis linking photosensitive skin injury to systemic organ damage, identifying two complementary therapeutic checkpoints.

The translational relevance of this pathogenic axis is highlighted by the availability of the FDA-approved CXCR4 antagonist AMD3100/plerixafor, together with PKCα-targeting agents, such as the pan-PKC inhibitor darovasertib or enzastaurin, a selective PKCα/β inhibitor currently in clinical trials (*60*). Because both PKCα and PKCβII are implicated in lamin B phosphorylation during nuclear lamina disassembly in other contexts (*13, 61, 62*), enzastaurin represents a promising candidate for modulating NETosis through nuclear envelope regulation. However, the specific contribution of PKCβII to NETosis requires further investigation. Whether these agents suppress UVB-triggered NETosis in patients with SLE remains to be determined in future clinical studies. Although NETosis has long been implicated in chronic lupus pathogenesis, the present study provides in vivo evidence that neutrophil NETosis is a critical initiating mechanism underlying acute, UVB-induced skin-kidney inflammation. By coupling localized skin injury into systemic kidney inflammation through neutrophil dissemination, this work defines a mechanistically distinct pathway for acute, photosensitivity-induced lupus exacerbation. Future studies should determine whether recurrent activation of this pathway contributes to chronic lupus progression and lupus nephritis.

## METHODS

### Sex as a biological variable

Given the profound female bias in human lupus (approximately 90% patients) and the accelerated disease kinetics in female MRL/*lpr* lupus-prone mice, all experiments within the lupus context exclusively utilized female mice. However, for *in vivo* neutrophil dissemination experiments using Cxcr4-deficient strains on a non-lupus background, both sexes were included.

### Mice

MRL/*lpr* lupus-prone mice and *Pkcα*-deficient mice (B6;129-*Prkca* ^tm1Jmk^/J mice, JAX #009068) (*63*), along with their corresponding littermate controls, were purchased from the Jackson Laboratory, and housed in a pathogen-free environment, and had *ad libitum* access to food and water. To generate the MRL/*lpr;Pkcα*^-/-^ strain, lupus-prone mice with *Pkcα* deficiency, *Pkcα*^-/-^ mice were backcrossed onto the MRL/*lpr* genetic background for at least 10 generations. For *in vivo* spatiotemporal tracking of pathogenic neutrophil dissemination from the skin to the kidneys, we generated *kikGR;Cxcr4*^+/-^ (Kikume Green-Red) mice, as homozygous *Cxcr4* deficiency results in perinatal lethality. Briefly, *Cxcr4^flox^*mice (JAX, #008767) were crossed with *CMV-Cre* mice (JAX, #006054) to create *Cxcr4*-deficient mice, which were then crossed with *kikGR*-expressing mice (JAX, #035393) (*44*). This model enabled the visualization of neutrophil trafficking via kikGR fluorescence, with *kikGR;Cxcr4^flox^*mice serving as controls. All animal experiments were approved by the Institutional Animal Care and Use Committee (IACUC) of the Philadelphia VA Medical Center.

### UVB exposure and sample harvest

The dorsal skins of 8-week-old, asymptomatic young female MRL/*lpr* or MRL/*lpr;Pkcα*^-/-^ mice were subjected to UVB irradiation (150 mJ/cm^2^/day) or sham treatment for 5 consecutive days under anesthesia, as previously described (*13, 14*). The selected UVB dose (150 mJ/cm^2^) has been widely used in murine models to reliably induce cutaneous inflammatory and immune responses (*64–68*). In separate pharmacological studies, eight-week-old asymptomatic female MRL/*lpr* mice received daily intraperitoneal injections of either the CXCR4 selective antagonist IT1t (10 mg/kg/day) or sterile normal saline (vehicle control). This dose was selected based on previous findings in another lupus mouse model (*59*). Pharmacological treatment began one day prior to the 5-day UVB regimen and continued for 6 consecutive days. Each experimental cohort consisted of 5-10 mice.

To enable the spatiotemporal tracking of neutrophil migration *in vivo* from the skin and their infiltration to the kidneys, *kikGR;Cxcr4*^+/-^ and *kikGR;Cxcr4^flox^* control mice were subjected to a sequential UVB irradiation and violet light photoconversion over five consecutive days. Each day, the shaved dorsal skin was first exposed to UVB irradiation (150 mJ/cm^2^/day (*13, 14*)). Then, the kikGR green protein expressed by infiltrating neutrophils was photoconverted to red fluorescence by applying violet light (405 nm), as described before (*44*). The violet-blue light was supplied by a 50 W LED lamp (0.37 numerical aperture fiber, Mightex) at 100 mW, illuminating round skin spots (∼2cm in diameter) on the marked area of UVB-irradiated dorsal skin for approximately 15 min per exposure (*33*). This sequential exposure converts the kikGR protein from green to red fluorescence, enabling the *in vivo* spatiotemporal tracking of UVB-activated neutrophils as they migrate from the skin to the kidneys.

One day after the last UVB irradiation, all mice were anesthetized and the systemic circulation perfused with EDTA-PBS (50 units/ml) via left-sided cardiac puncture (*69*). Then the perfused kidneys and dorsal skin were harvested for subsequent experiments. For histologic studies, the skin or kidney tissues from all mice were fixed in 4% paraformaldehyde (PFA) for 24h, embedded in paraffin, and sectioned at a thickness of 5 µm (*13, 14*). Proteinuria score was measured for all mice before sacrifice using Chemstrip 2GP (Roche, Indianapolis, IN). A scale of 0–3 was used to correspond to the following total urinary protein levels: 0 (negative), 5–20 mg/dL (trace), 30 mg/dL (+), 100 mg/dL (++), and 500 mg/dL (+++).

### Cell Culture and Treatment

Mouse bone marrow (BM) neutrophils from the *femur* and *tibia* BM of MRL/*lpr* mice were isolated as described in our previous publications (*13, 14*), and were purified with a neutrophil isolation kit (Miltenyi Biotec). All of the above primary mouse neutrophils were treated without or with 5 or 10 µM PAF, or irradiated without or with UVB at 30-60 mJ/cm^2^, and cultured for 20h (*13, 14*) for analysis of neutrophil expression of IFNγ and IL-17A cytokines, complement C3 (C3), or CXCR4. For induction of IFNα in neutrophils, these neutrophils were co-incubated with purified DNA-NETs that were prepared and isolated as described before (*13, 14*). Then, neutrophils were fixed with 2% PFA and stored at -20 °C for later analysis.

### Flow cytometry analysis

To detect UVB-induced neutrophil expression of CXCR4, IFNα, IFNγ, IL-17A, and C3, primary MRL/*lpr* mouse neutrophils were treated without or with PAF (10 µM PAF) or UVB (60 mJ/cm^2^), and incubated for 20h, followed by fixation with 2% PFA. Then these cells were first permeabilized with 0.1% Tween-20 and co-stained with PE-labeled anti-mouse Ly6G, FITC-labeled anti-mouse IFNα, IFNγ, IL-17A, or C3, and APC-labeled anti-mouse CXCR4. Analysis of expression of CXCR4, and/or cytokines, C3 in neutrophils were based on gated cells that were Ly6G positive cells. Flow cytometry analyses were conducted by FACS Canto, and data were analyzed by Flowjo software as described before (*14*).

### Transmigration Assays

To study the effects of CXCR4-CXCL12 signaling pathway on neutrophil migration in the context of UVB, the transmigration ability of primary neutrophils from MRL/*lpr* mice was examined by using Boyden chamber as previously described (*13, 14, 42*). Primary neutrophils from MRL/*lpr* mice were first stimulated by 10 µM PAF for 20h to induce their CXCR4 expression. To study the effects of CXCR4 inhibitor, neutrophils (4 x 10^5^) from MRL/*lpr* mice with PAF pretreatment were treated or not by 10 µM IT1t, a selective CXCR4 antagonist, for 30 min before adding to the upper chamber with 3 µm pore inserts of 24-well trans-well plates, followed by stimulation with CXCL12 (300 ng/ml) in the bottom well for 5h. The pore transmigration ability was expressed as relative transmigration compared to that in control neutrophils of the mice without treatment as described before (*14*).

### Immunohistochemistry analyses of the skin and kidneys

The paraffin-embedded skin or kidney tissue sections of all mice were stained either by hematoxylin and eosin (H&E) or by fluorescent staining as described before (*13, 14*). Infiltration of inflammatory cells in skin of all mice was counted for the nucleated cells in the epidermis, dermis, or fat of the H&E stained skin sections as described in our published works (*13, 14*). The glomerular cellularity, including podocytes, endothelial cells, mesangial cells, and infiltrated cells of the kidney cross sections of all mice, was counted as described before with modifications (*70*). In fluorescent staining, the neutrophil markers PE-labeled anti-ly6G or Alexa-647-labeled myeloperoxidase (MPO) were used for detection of neutrophils and NETosis in combination with FITC-labeled citH3, and DNA staining with DAPI as described before (*13*). FITC-labeled anti-mouse antibodies against IFNα, IFNγ, IL-17A, C3, or CXCR4 were used to stain the corresponding molecules in the skin or kidneys. The fluorescently matched isotype control immunoglobulins were used as isotype control staining. Slides were mounted with Gold Antifade Mountant (Invitrogen).

The fluorescent images were analyzed with an Olympus Fluoview 1000 confocal microscope or Leica fluorescent microscope. To analyze images for molecular quantification in IHC analyses, the 2D high resolution fluorescent images from at least 5-6 random non-adjacent areas of dermis per skin section, or 20 random glomeruli per mouse kidney section, were captured with an Olympus confocal or Leica fluorescent microscopes and saved as .tiff files. The images were analyzed with NIH ImageJ software by following step-by-step workflow according to the published protocols (*71, 72*). Quantification of the molecules of interest, i.e. neutrophils, NETs, cytokines, or C3, and CXCR4 were based on an automatic threshold analysis of immunofluorescence images to automatically identify the top brightest structures of each image as described in our published work (*13, 14*).

### Statistical analyses

Statistical analyses were performed using GraphPad Prism 6. Normally distributed data are shown as means ± standard deviations (SD). Comparisons between two groups were conducted with Student’s t test. Comparisons among three or more groups were performed using ANOVA, followed by Student-Newman-Keuls (SNK) test. Pearson’s correlation coefficient was used for analyses of a linear association between two variables. Statistical significance was considered at a level of *P*-value < 0.05.

## AUTHOR CONTRIBUTIONS

The research was conducted at the University of Pennsylvania and VAMC in Philadelphia. Liu conceived and designed the study. Lyu, Li, Yi, and Liu performed the laboratory experiments and acquired the experimental data. For the microscopic image analysis, Lyu, Liu, Yi took the images, Lyu and Li conducted further quantification and statistical analysis. Liu and Lyu analyzed and interpreted the data and finalized the paper. Lyu, Li, Zhang, Wei, Werth and Liu were involved in the investigation, preparation and revision of the manuscript. All authors read and approved the final manuscript.

## ACKNOWLEDGEMENTS

The authors would like to acknowledge Penn Skin Biology and Diseases Resource-based Center for their skin histology service. The authors would thank Debra A. Pawlowski (animal facility, Philadelphia VA Medical Center) for her kindly help with our animal experiments. This work was supported by Lupus Research Alliance (416805) and NIH R21AI144838 (to MLL), Veterans Affairs Merit Review Award (1I01BX005921) and NIH R01 5R01AR076766 (to VPW), NIH 1R41AR082749-01 (to Werth, Liu).

## Supplemental Materials

### Supplemental Data

**Suppl Fig 1.**
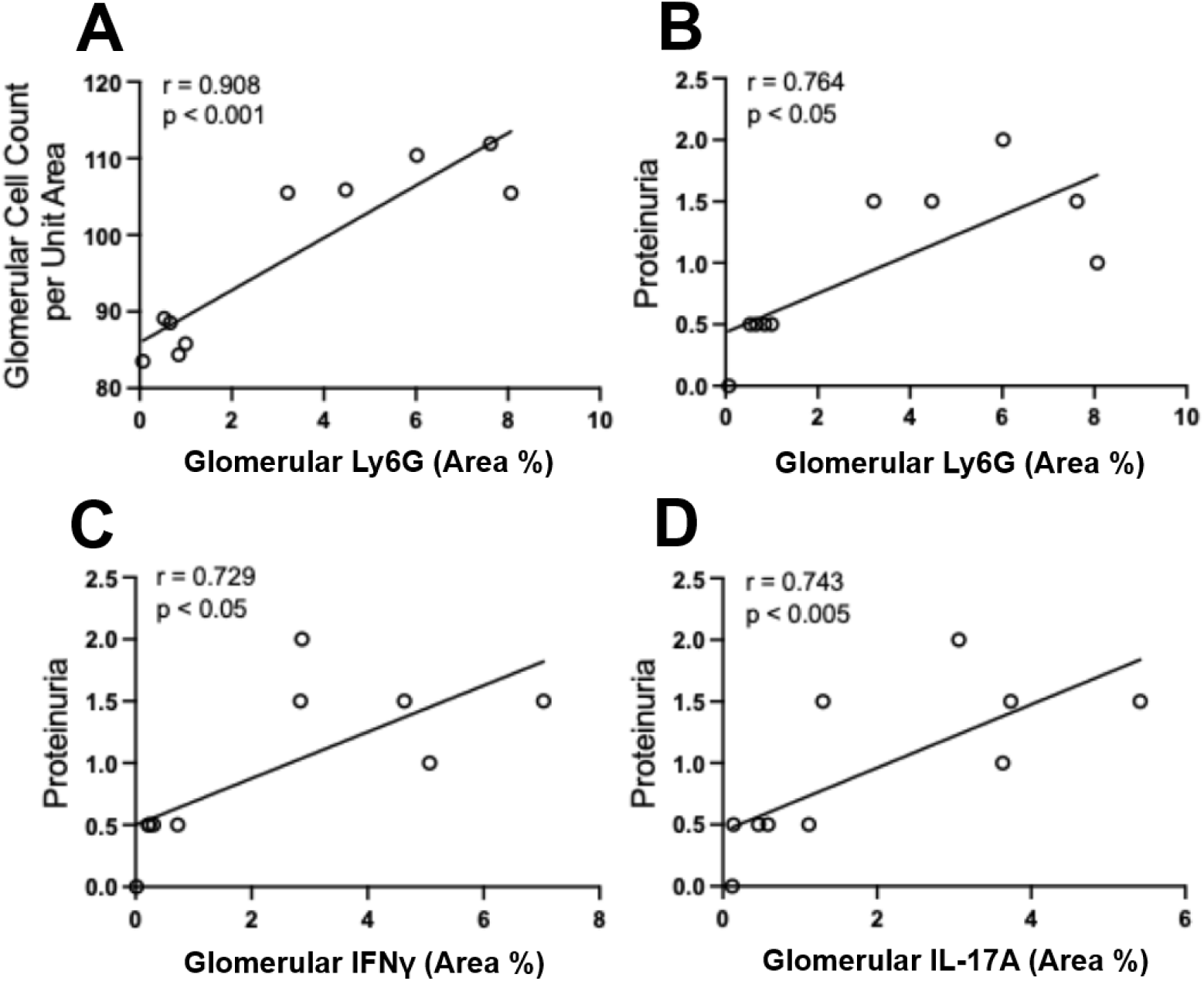
Renal neutrophil infiltration and NET-associated cytokines correlate with proteinuria in acute kidney inflammation in lupus-prone mice triggered by UVB. **A.** Correlation between glomerular neutrophil infiltration and glomerular cellularity. **B-D**. Correlation between glomerular neutrophils, NET-associated IFNγ, IL-17A with proteinuria in combined experimental groups. All data were analyzed using Pearson correlation coefficients.

**Suppl Fig 2.**
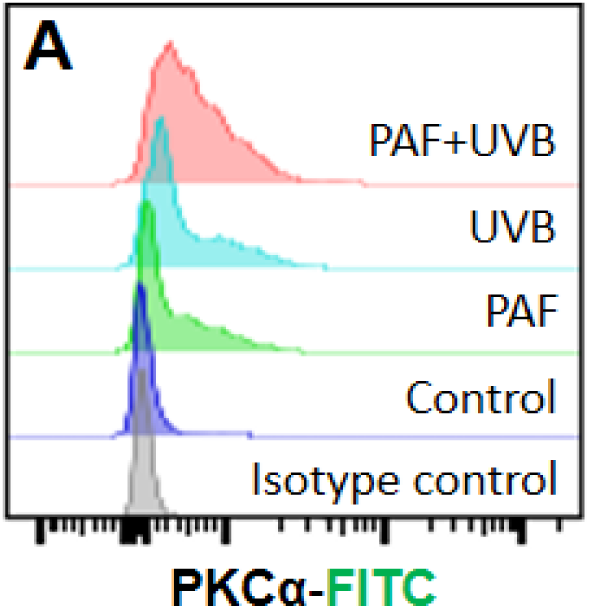
UVB induced expression of PKCα *in vitro* in primary neutrophils from lupus-prone mice. **A.** Representative flow cytometry histogram of PKCα expression in primary neutrophils from lupus-prone mice. Cells were stimulated without (control) or with 10 µM PAF, or 60 mJ/cm^2^ UVB irradiation, or both for 20h, then processed and immuno-stained with FITC-labeled anti-mouse PKCα. All neutrophils were gated on ly6G^+^ cells.

**Suppl Fig 3.**
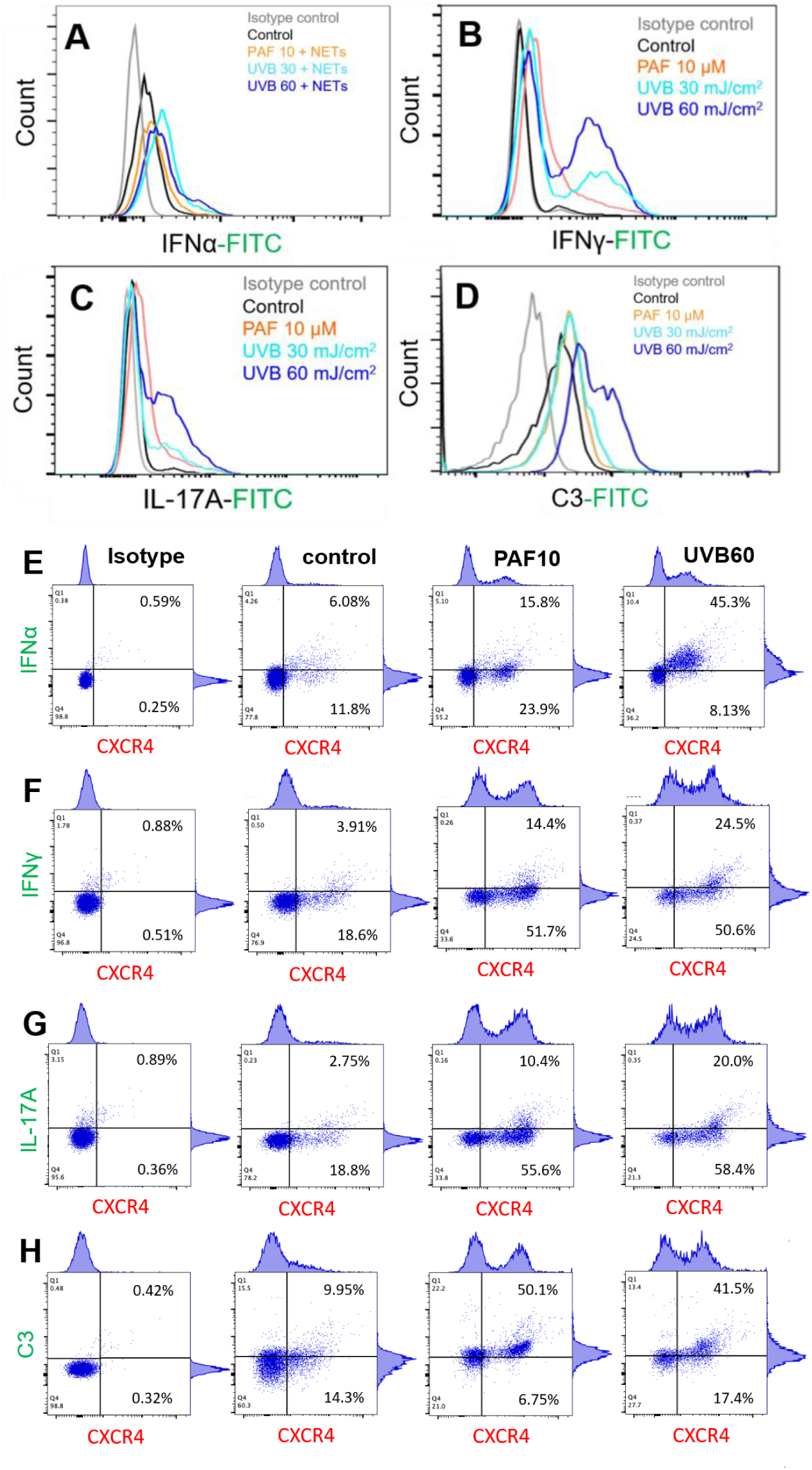
UVB induced expression of cytokines and C3 *in vitro* in primary neutrophils from lupus-prone mice. **A-D.** Representative flow cytometry histogram of IFNα, IFNγ, IL-17A, or C3 expression in primary neutrophils from lupus-prone mice. Cells were stimulated without (control) or with 10 µM PAF, or 30-60 mJ/cm^2^ UVB irradiation, and incubated for 20h, then processed and immunostained as detailed in methods and Fig 5. **E-H**. Representative flow cytometry dot plots of CXCR4 co-expression with IFNα (E, DNA-NETs were presented during stimulation), IFNγ (F), IL-17A (G), or C3 (H) in primary neutrophils from MRL/lpr mice, which were stimulated without (control) or with 10 µM PAF or 60 mJ/cm^2^ UVB for 20h. All neutrophils were gated on ly6G^+^ cells.

**Suppl Fig 4.**
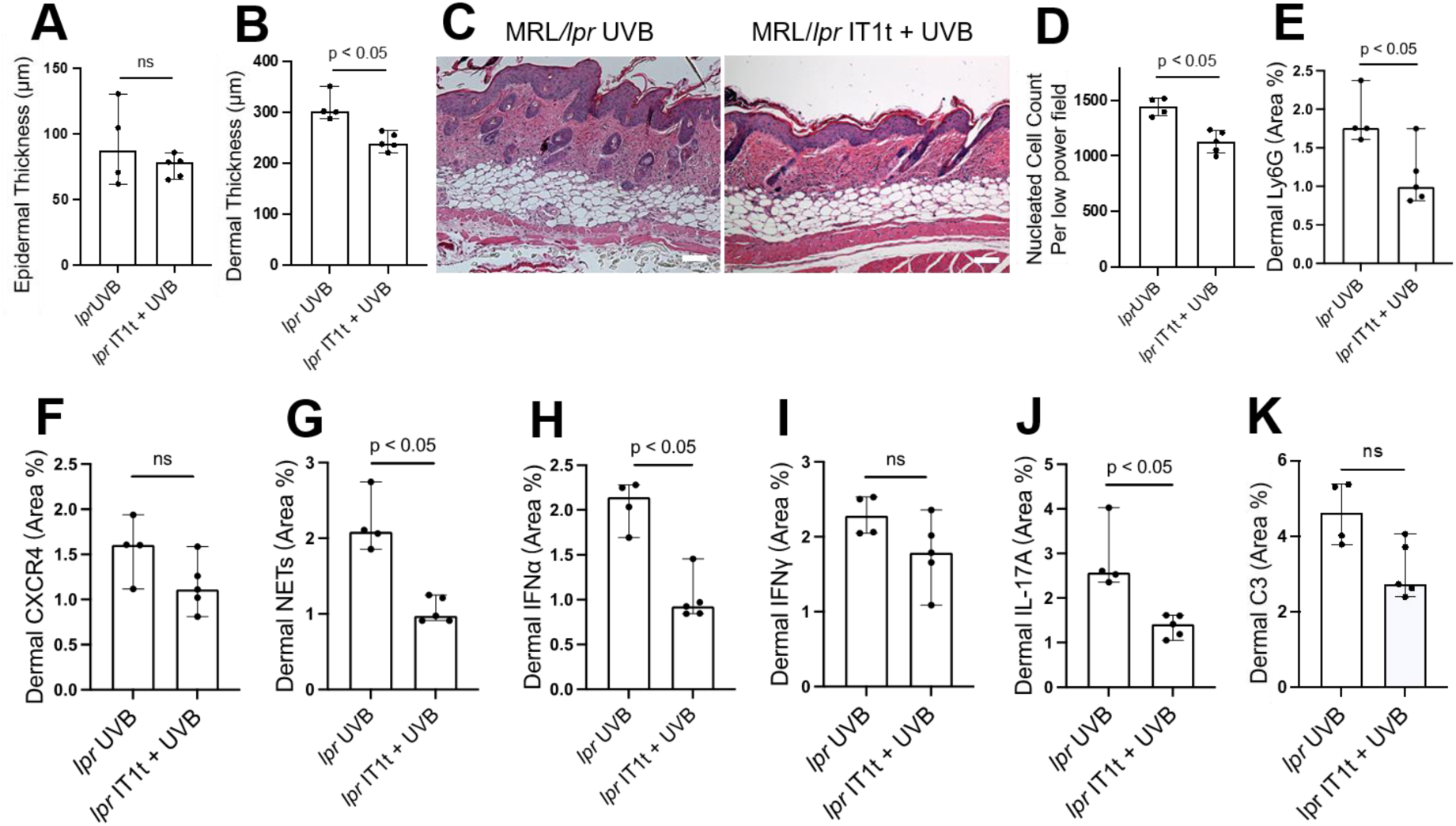
Effects of pharmacological CXCR4 inhibition on acute skin inflammation in UVB-irradiated lupus-prone mice. **A-K**. Summary analyses or representative H&E images of epidermal (A) and dermal (B) thickness, inflammatory cell infiltration (C-D, nucleated cells), neutrophils (E), CXCR4-expressing neutrophils (F), NETs (G), and NET-associated IFNα (H), IFNγ (I), IL-17A (J), or C3 (K) in skin of UVB-irradiated MRL/*lpr* mice without (lpr UVB) or with IT1t administration (lpr IT1t+UVB). Scale bars, 100 µm. Results represent 4-5 biological replicates. Panels A,B,D-K display means ± SD determined by Student’s t test for two-group comparisons.

